# 3D Digitizing An Animal Organ Could Make It Possible To Work With Anatomy Classes In The Metaverse Or With A 3D-Printed Real-Scale Organ On Demand

**DOI:** 10.1101/2022.10.24.513584

**Authors:** Dogasav Sumer, Onat Ozten, Mert Gungor, Meryem Yilmaz, Ismail Nart, Cihan Tastan

**Author notes:** Equally contributed first authors (Dogasav Sumer, Onat Ozten, Mert Gungor). **Author Contributions:** D.S., O.O., M.G., and C.T. designed, completed, and/or analyzed experiments. M.Y. and I.N. contributed to 3D printing. D.S., O.O., M.G., and C.T. wrote the manuscript and contributed to the editing of the manuscript. C.T. led the project and contributed to the design and interpretation of the data.

## Abstract

Organ scanning is not a novel technology but the unique part of this study is to eliminate the need for expensive devices by using free iOS/Android applications. The apps allow users to create high-quality 3D models from photos taken with a phone or tablet. Metaverse and 3D digitization are the future of technology. Implementation of these approaches into the anatomy can change education style. Also, 3D printing technology along with the digitization of real-sized organs enables researchers and clinicians to study with 3D printed organs in class, even with augmented reality glasses. In this study, we first aimed to 3D digitized an animal organ with its real sizes to print in a 3D printer. Next, we wanted to digitize decellularized form of the organ, which suggests researchers study extracellular matrix and vascularization. This digitization of the different organ forms can improve the progression of artificial organ studies. Free and easy-to-use iOS or Android applications for 3D digitization enable medical students to study not only animal organs but cadavers. Hereby, we are suggesting organ scanning can be done on a daily basis in a standard laboratory. It is aimed to enable medical students to work effectively in anatomy lessons in Metaverse or in real cases, thanks to the anatomical library created with the accessibility brought by easy usability.

## Introduction

Organ scanning is not a novel technology but the unique part of this study is to eliminate the need for expensive devices by using iOS or Android phones and applications (Apps). The 3D capture apps allow users to create high-quality 3D models from photos taken with a phone or tablet. Biologic scaffolds derived from decellularized tissues and organs have been used successfully in pre-clinical animal studies as well as human clinical trials. When cells are removed from a tissue or organ, the complex mixture of structural and functional proteins that comprise the extracellular matrix remains (ECM) [1]. Decellularized organs are ideal transplantable scaffolds because they contain all of the necessary microstructure and extracellular cues for cell attachment, differentiation, vascularization, and function [2]. The intention of scanning ECM and bioprinting it is for further recellularization studies based on stem cell research [3]. Bioprinted ECM provides an optimal environment for cells to grow and differentiate.

Hereby, we are suggesting organ scanning can be done on a daily basis in a standard laboratory. It is aimed to enable medical students to work effectively in anatomy lessons thanks to the anatomical library created with the accessibility brought by easy usability. After the development of three-dimensional bioprinters, the idea of printing an organ and transplanting the said organ to a living organism has been on the minds of scientists. In light of this aspect, we designed an experimental procedure consisting of several stages; the first one is the digitalization of the sheep heart the second decellularization of sheep heart tissue and obtaining the extracellular matrix (ECM), and finally printing the digitalized ECM from 3D printers. This study enables medical students to work effectively in anatomy lessons in Metaverse or in real cases with 3D-printed true-scaled organs.

## Material

### 3D Scanning

8-12 months old sheep hearts were obtained from slaughterhouses. The sizes of the organs were measured with the L ruler. For 3D scanning of the organs, a platform was created where the newly bought sheep heart can rotate 360 degrees freely. Then, an average of 130 photos were taken with the Polycam application with a mobile device (iPhone 12 used) to scan all parts of the heart, and the scanning process was performed. Afterward, the 3D version of the photos on the device was brought together and transferred to the computer environment to be printed on the 3D printer. The suitability of the original heart dimensions was tested and the printout was taken from the 3D printer.

### Decellularization

To decellularize the hearts, they were kept at −80 degrees for 20 minutes, then thawed at 37 degrees for 20 minutes in a shaking incubator. This process was performed 4 times. Following washing the heart 6-8 times with PBS, with a syringe for the vessels, then washed with 50 mL of deionized water. Next, gently washed with 50 mL of 1% SDS. The organs were aged/incubated for 96 hours in 150 mL 1% SDS on a shaker. The hearts were kept overnight without SDS, washed in 50 mL of PBS, and incubated in 150 mL of PBS for 24 hours.

## Result

In the experiment, we aimed to 3D digitize an animal heart before printing **(Fig. 1A and 1B).** Next, we tested the 3D printing of the true-sized organ **(Fig. 1B).** Before 3D printing studies in order to see the accuracy of the scanning procedure and a free iOS/Android application and a standard 3D printer were used. To prove the scanning of decellularized heart, we performed freeze-shock and SDS/PBS incubation of the organ **(Fig. 1C).** The main reason of this study is to obtain the ECM structure of the heart and scan the decellularized ECM.

**Figure 1:**
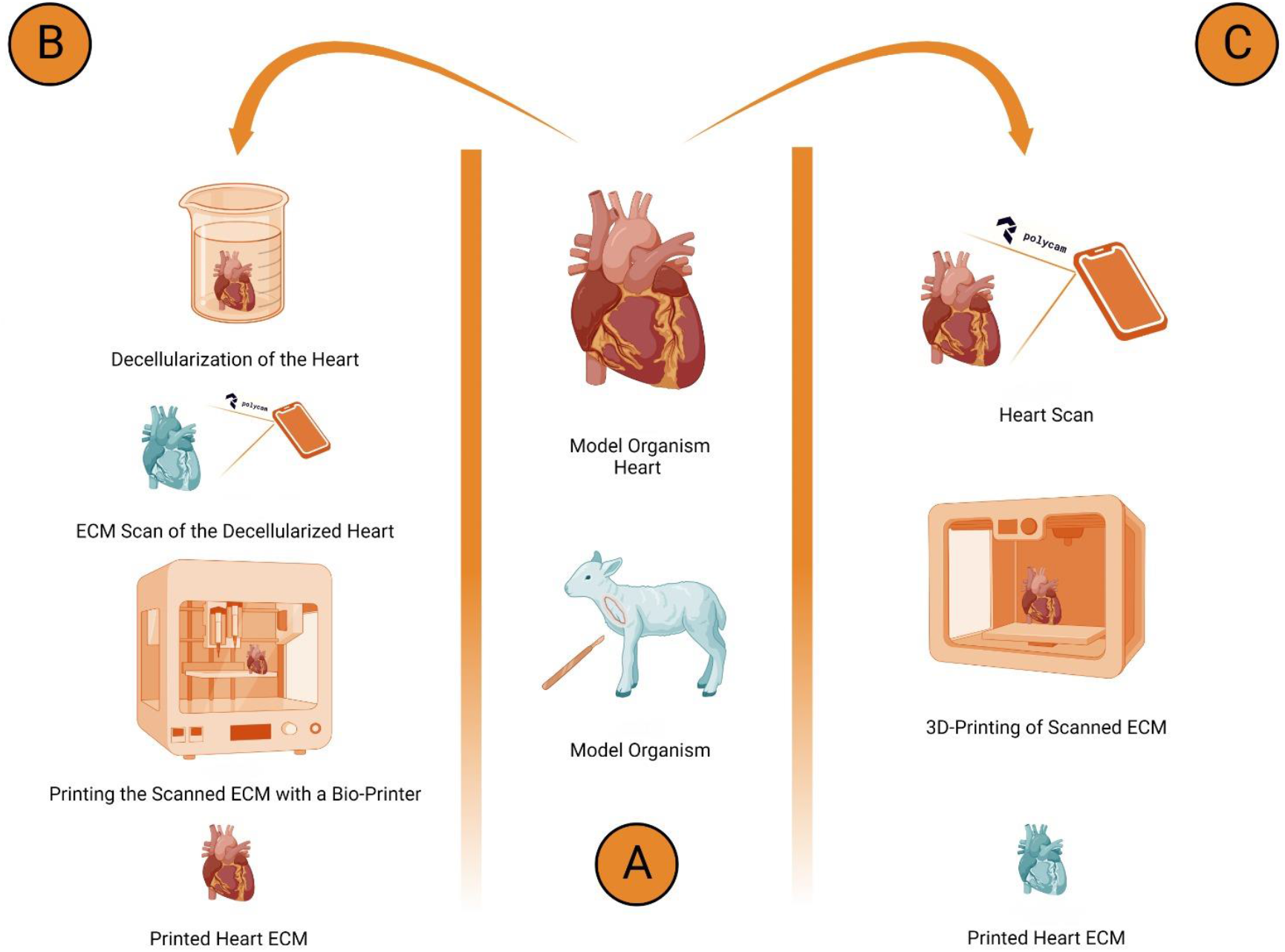
An experimental setup of the study with 3D digitization, decellularization, and 3D printing of a model organism, sheep, and heart. **A**. Process of organ extraction from a model organism, sheep, is shown. **B.** The scanning process with an iOS/Android app, Polycam, and standard 3D printing of the scanned heart. **C**. Decellularization process of the heart and scanning of the processed organ’s ECM.

To perform this setup, hearts were measured and taken after slaughter. The Y axis (length) is measured as 10.9 cm and the X axis as 5.9 cm, as a mean of three sheep hearts **(Fig. 2A).** The measurements were confirmed by rechecking the heart after being digitized and transferred to the 3D printer **(Fig. 2B).** The 3-dimensional state of the heart taken from our animal model has been transferred to the virtual environment, almost at the same level as the real one **(Fig. 2C).** You can refer to the appendix for a detailed review. In the setup we have arranged, the actual structure and size of the heart have been digitized without any pressure.

**Figure 2:**
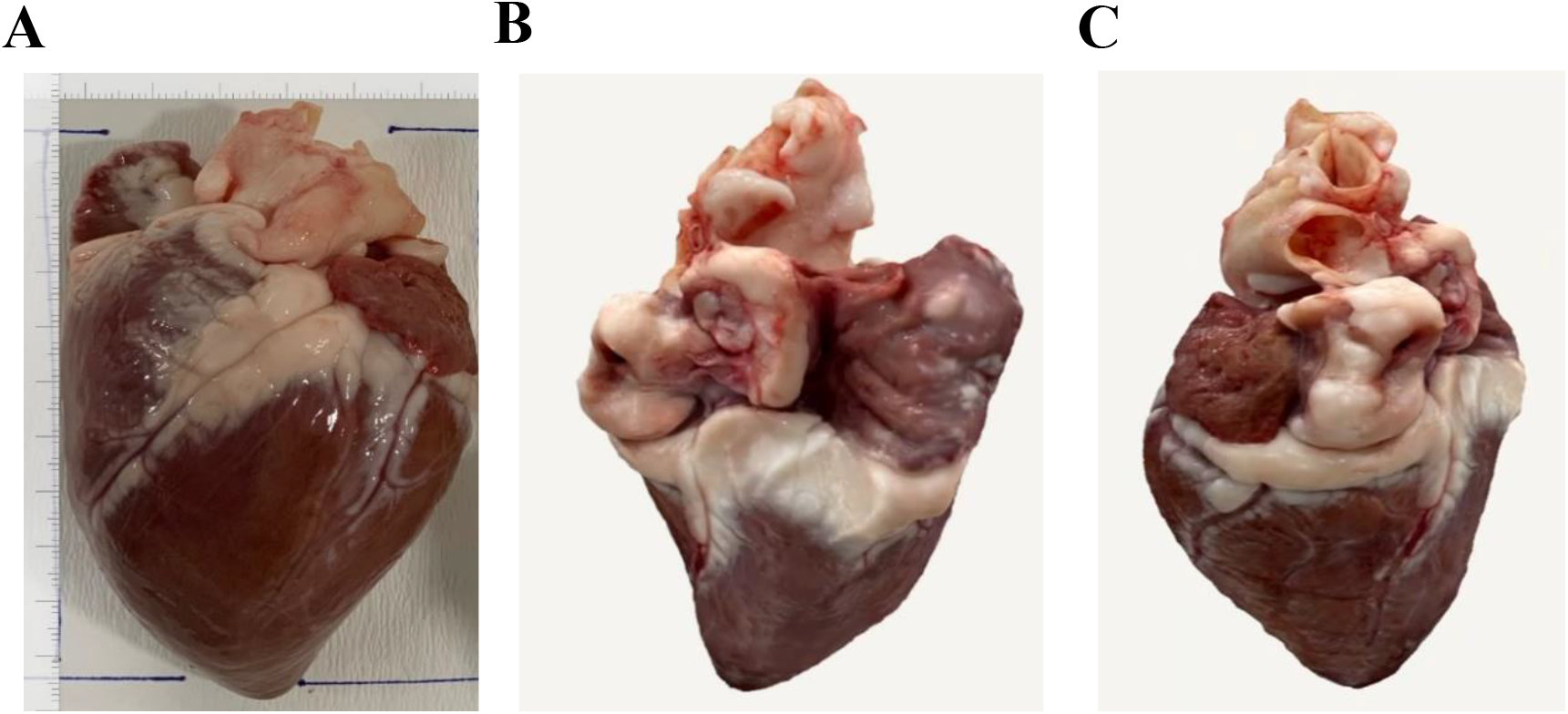
3D digitization of the sheep heart. **A.** The picture showing measurements of the hearts (mean of the measurements,10.9 cm on the Y axis, 5.9 cm on the X axis) **B.** and **C.** Digitized version of the hearts using the app.

Next, we wanted to 3D print the true-sized organ with a conventional 3D printer. The aortic vascular structure, right and left atrium parts, and ventricles found in a normal heart were printed at such a high level of detail that they could be observed in the printed heart **(Fig. 3)**.

**Figure 3.**
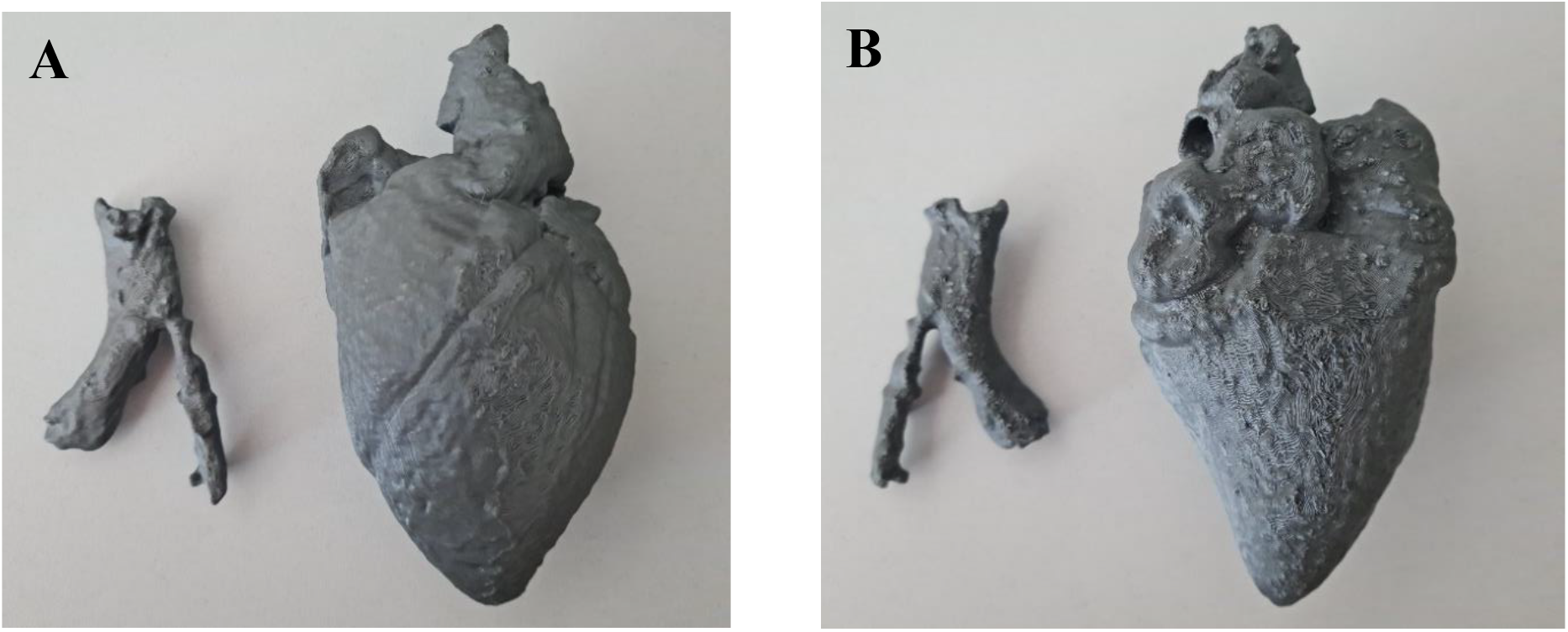
3D printed form of digitized true-scaled sheep heart and aort.

Then, we asked to digitize a decellularized heart to visualize vascularization and ECM after trypan blue staining of the veins. The decellularized and 3D digitized heart were made to appear more clearly **(Fig. 4).** Contrary to normal decellularization experiments, in our process, since it is not processed with the pump assembly, a complete decellularization process has not been realized. The decellularization process is seen more clearly in the heart in the places indicated by the arrows, that is, in the parts close to the main vessels. This test showed that decellularization of the organs along with 3D digitization may help anatomy studies.

**Figure 4:**
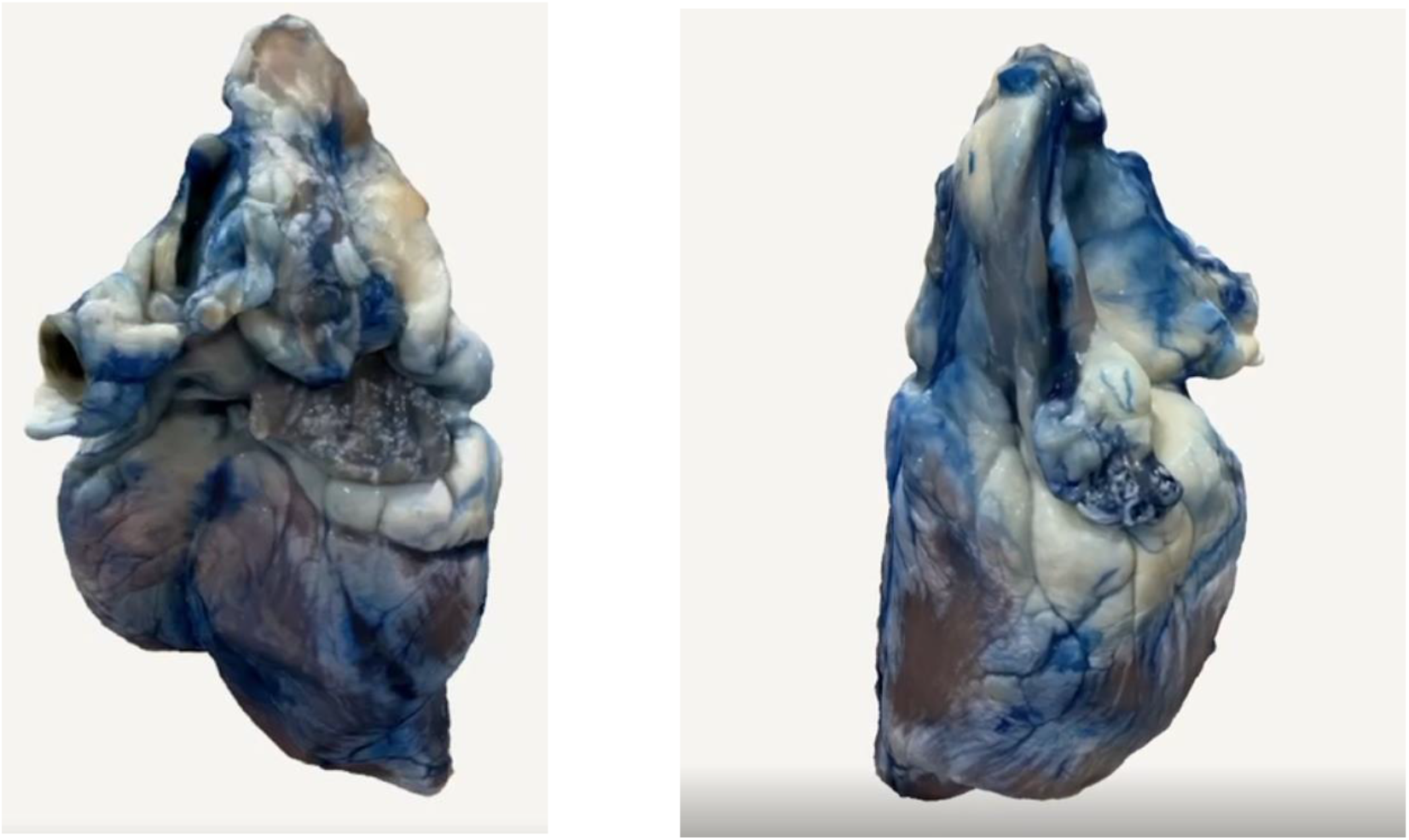
3D digitized 2-week decellularized sheep heart stained with trypan blue.

## Discussion

3D scanning is not a novel technology but the unique part of this study is to eliminate the need for expensive devices by using free iOS/Android applications. The apps allow users to create high-quality and real-scaled 3D organ models digitized from photos taken with a phone or tablet. Metaverse and 3D digitization are the future of technology. Implementation of these approaches into the anatomy can change education style. Also, 3D printing technology along with the digitization of real-sized organs enables researchers and clinicians to study with 3D printed organs in class, even with augmented reality glasses. In this study, we first aimed to 3D digitized an animal organ with its real sizes to print in a 3D printer. The easy use and accessibility of the application and scanning technology we use will enable everyone to scan without the need for any special training. This will serve as a facilitator in anatomy lessons and research of medical and biotechnology students in anatomy libraries to be established in the future.

Next, we wanted to digitize decellularized form of the organ, which suggests researchers study extracellular matrix and vascularization. Due to the changes we made within the scope of laboratory facilities, a homogeneous decellularization did not appear as in the Delgado at al. article we reviewed [3]. The problem can be solved with the appropriate pumps and chemicals in the article. In addition, in this experiment, with the needling technique for the decellularization process, the SDS chemical was passed into the vessels more and accelerated the process. Despite this change, it was shaken continuously at 56 rpm on a shaker for 2 weeks in the absence of pump and necessary chemicals. The SDS page we used is the ‘Merck’ branded SDS prepared for the running gel. We think that since there is a different material than SDS which is Edetate disodium dihydrate, it is not fully penetrated into the cells.

Easy-to-use iOS or Android applications for 3D digitization enable researchers to study organs and cadavers. Hereby, we showed organ scanning can be done on a daily basis in a standard laboratory to enable medical students to work effectively in anatomy lessons in Metaverse or in real cases.

